# Secondary Motor Cortex Transforms Spatial Information into Planned Action During Navigation

**DOI:** 10.1101/776765

**Authors:** Jacob M. Olson, Jamie Li, Sarah E. Montgomery, Douglas A. Nitz

**Affiliations:** Brandeis University; Stanford University; Icahn School of Medicine at Mount Sinai; University of California, San Diego

## Abstract

Fluid navigation requires constant updating of planned movements to adapt to evolving obstacles and goals. A neural substrate for navigation demands spatial and environmental information and the ability to effect actions through efferents. Secondary motor cortex is a prime candidate for this role given its interconnectivity with association cortices that encode spatial relationships and its projection to primary motor cortex. Here we report that secondary motor cortex neurons robustly encode both planned and current left/right turning actions across multiple turn locations in a multi-route navigational task. Comparisons within a common statistical framework reveal that secondary motor cortex neurons differentiate contextual factors including environmental position, route, action sequence, orientation, and choice availability. Despite significant modulation by context, action planning and execution are the dominant output signals of secondary motor cortex neurons. These results identify secondary motor cortex as a structure integrating environmental context toward the updating of planned movements.

## Introduction

Navigation is a cognitive process necessitating neural encoding of complex spatial relationships between an organism and its environment. This information must then be used to generate specific action plans. Often, obstacles and threats or rewards exist such as unpassable terrain, predators, food, or social opportunities. Under these circumstances, multiple features of spatial context must be integrated in order to define the appropriate action to take, creating incentives for complete representations of environmental features. Finally, fast, uninterrupted movement to a goal also demands that upcoming actions be planned in advance of their execution, but as most environments are filled with ever-changing obstacles, barriers, and even goals, this process must also be capable of being dynamically updated.

A large body of research describes how sensory information related to self-motion, boundaries, and landmarks is transformed, through neural circuitry, to help form a cognitive map. Sensory information, initially encoded in egocentric reference frames^1-5^, is converted into representations of position and orientation relative to environmental boundaries and landmarks^6-15^. The latter forms of information are described by a frame of reference largely independent of the animal itself – an allocentric frame of reference. Together, they form a distributed network serving to synchronize the organism’s internal representations to the environment.

However, spatial cognition must ultimately be transformed into a motor output. Specific features of cortico-hippocampal connectivity point to retrosplenial and posterior parietal cortex as intermediaries in this encoding process. Retrosplenial (RSP) and posterior parietal (PPC) cortices are interconnected with hippocampus, subiculum, and perirhinal, postrhinal, and entorhinal cortices^16-24^. These two regions contain neurons representing information in multiple egocentric and allocentric reference frames^9, 25-33^ and boast dense projections to secondary motor cortex (M2)^34-36^. In turn, M2 projects strongly to primary motor cortex and brainstem and spinal regions involved in motor control^37,38^ and could therefore be a structure important to the transition of integrated spatial information into action as part of the larger navigational process^35,39^.

M2 has been the subject of much experimental work aimed at determining critical neurophysiological components of decision-making processes, rule implementation, and action planning and execution as instructed by single modality sensory cues^40-47^. Anatomically, M2 in the rat is a subregion of prefrontal cortex that, historically, has been referred to by a number of different names including: shoulder region, anterolateral motor cortex, medial precentral cortex, Fr2, medial agranular cortex, premotor cortex, dorsomedial prefrontal cortex, and frontal orienting field. With respect to navigation, published work indicates that M2 neuron firing predicts upcoming navigational choices from a single environmental location^41^ and that neighboring prefrontal cortex sub-regions encode information related to rules for navigation as well as anticipated effort and reward associated with a route^48-62^. Yet, whether M2 neuron firing dynamics are consistent with a role transitioning a broader and spatially distributed set of organism-environment spatial relationships into planned actions remains largely an open question. The answer has critical importance for developing systems neuroscience models of the navigational process.

In the present work, we examined M2 neural activity patterns in the context of a large ‘triple-T’ maze. Rats repeatedly traversed specific routes to and from multiple goal locations. The chosen environment and task structure spatially and temporally distanced individual actions from surrounding actions and goal locations. This enabled evaluation of distinctions in firing associated with action planning versus execution as well as the influence of multiple spatial variables.

We find that M2 neurons most robustly discriminate left versus right turning actions and do so reliably across all turns. Across M2 neurons, activity differentiating turn type is most often maximal near the peak of a turn, but, for many neurons, peak firing occurs well before the action is actually executed. While holding action type constant, M2 neuron populations were also found to represent multiple spatial and directional features that varied across turn locations. Such encoding took the form of reliable modulation in the intensity of turn-related spiking activity, demonstrating that M2 ensembles simultaneously encode current and planned actions as well as the navigation-relevant context of those actions. M2 neuron firing also varied according to the presence or absence of choice at a turn, suggesting a more complex role in the action preparation process than simple categorical left/right choice. Together these findings implicate M2 as a structure capable of transforming spatial, directional, and decision-making context into the actions that define navigational behavior.

## Results

Rats were trained on a triple ‘T’ track maze (Figure 1A) to traverse eight three-turn ‘internal’ routes of the same total length. All eight internal routes shared a single start location but led to eight different goal locations where food reward (1/2 piece Cheerios cereal) was delivered. Animals then returned to the start location via the pathways of their choice available along the maze perimeter. Animals were allowed to move freely in all areas except across the internal route spaces where backtracking was prevented. Correction was rarely necessary during recordings as animals were extensively trained to reach a criterion of 80% uninterrupted route traversals within individual sessions. The animals’ ability to traverse the routes uninterrupted persisted after surgical implantation of the recording headstage. The rats averaged 64% route traversals per recording session categorized as smooth, uninterrupted running (Figure 1B). Henceforth we will refer to these route traversals as “clean” runs. Animals averaged a velocity of 42cm/s during clean runs. Animals traversed routes according to one of three separate reward schedules, “high-low”, “visit-all-8”, or “visit-all-4” (see methods section for further detail). In the analyses to be presented, data from these different reward schedules were pooled.

**Figure 1:**
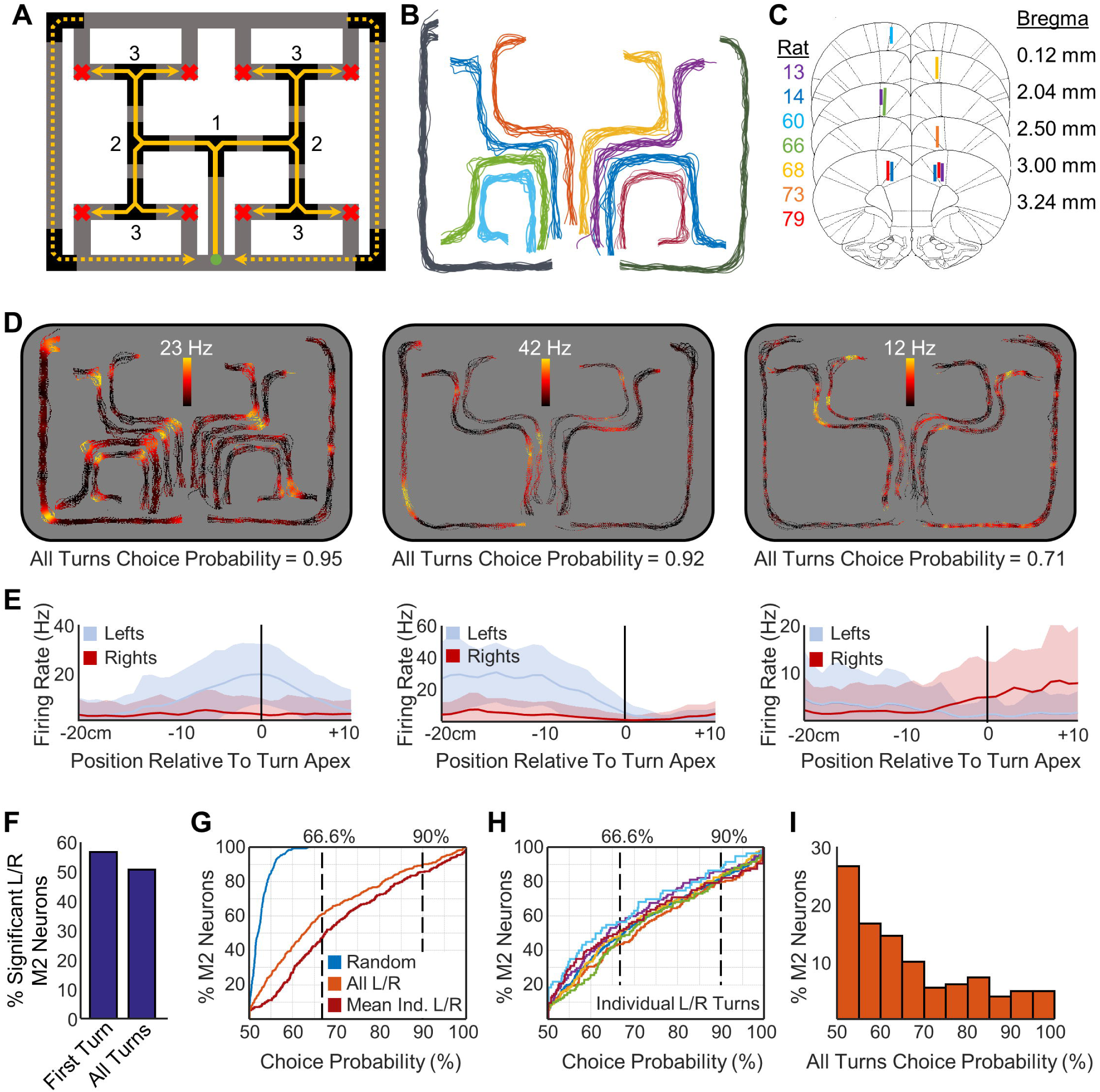
M2 robustly encodes action across space. **A)** Schematic of triple ‘T’ maze and route-running task. Animals made runs along each of up to eight partially overlapping internal routes (solid yellow lines) on the 160cm × 125cm track apparatus, leading from a start site (green circle) to any of four goal sites (red ‘x’). Numbers indicate the index of the turn in the progress of an internal route. From each goal site, the animal returned to the start via either of two return paths (dashed yellow lines). For some task setups, only the routes leading to the top four goal locations were used. **B)** Behavioral tracking of identified clean runs. Shown are all clean runs from one recording session in the visit-all-8 reward setup. Each color shows a separate route. Routes are minimally translated and stretched for visualization purposes. **C)** Electrode placement and recording ranges in M2 (seven rats). Lines track dorsal-ventral recording depths across recordings. **D)** Example neuron positional firing rate maps. Shown are the firing rates color-mapped as a function of route and track position. As in Figure 1B, routes are minimally translated and stretched from the actual track location to separate each map for visualization purposes. Under each rate map is the choice probability (CP) for left and right actions, pooled across all locations. **E)** Linearized perievent firing rate maps. Mean (line) and s.d. (shaded) of the individual linear firing rate values surrounding each of left and right turns, shown in blue and red, respectively, for the corresponding neurons in **D. F)** Percentage of M2 neurons significantly discriminating left/right actions for the first internal turn versus all turn locations combined. **G)** Cumulative density functions (CDFs) of the entire M2 populations’ all-turn-locations-combined left/right turn CPs (orange), mean action CPs of individual locations (maroon), and shuffled action CPs (blue). **H)** CDFs of the entire M2 populations’ action CPs of individual locations. Each color is a different spatial location on the maze. **I)** Probability density function of the action CPs for all turns pooled across locations.

Single unit activity was recorded from 73 left hemisphere and 230 right hemisphere M2 neurons (303 total) from seven rats recorded under the aforementioned conditions (Supplemental Table 1). Electrode tracks and endpoints are depicted in Figure 1C with example histological data in Supplemental Figure 1. To avoid biasing assessment of the population distribution of neural firing responses, all cleanly clustered neurons were included in analyses, regardless of firing rate. As such, the distribution of firing activity found is skewed towards lower rates consistent with a log normal distribution as described in other cortical regions^63^ (Supplemental Figure 2).

### M2 Robustly Encodes Actions across Contexts

Action encoding is prevalent in M2 across all turn locations and contexts. Firing rate maps of clean run traversals from three sample neurons are shown in Figure 1D. To quantify these findings, we defined a turn space around each track turn from 20cm before to 10cm after the turn apex (Figure 1E). This space was the maximal possible that prevented overlap with adjacent turn spaces. We required at least 8 clean runs through each grouping analyzed (mean number of clean runs = 32±14). Fifty-seven percent (168/296) of all M2 neurons had significantly different mean firing rates for the first internal path turn (Figure 1A, turn labeled 1, two-sided Mann-Whitney U test, α<0.05, n = 32±14 traversals).

In order for an empirically decoded action signal to be actionable, the information must be accessible to a downstream reader. Consistency in the action code despite variability in the actions’ contexts (e.g., environmental locations) would enable downstream motor cortex to decode actions without simultaneously decoding context. For example, if a neuron fires strongly for a specific action at one location, but lower at another, downstream neurons capable of driving execution of the action effectively receive no information concerning the intended action unless the animal’s location is concurrently transmitted. To truly encode an action itself, there must be a reliable action-encoding signal that dominates that of concurrent context.

Such reliability in encoding of left/right turning action was observed in M2 neuron populations. Action encoding was extremely consistent across turn locations and contexts as nearly as many neurons, 51% (154/303) of all M2 neurons, significantly discriminated turn direction even when the data was pooled across all turn locations (two-sided Mann-Whitney U test, α<0.05, n = 32±11 traversals, sample sizes matched to first turn sample sizes; 203/303 for two-sided Mann-Whitney U test α<0.05, n=131±52 traversals, full samples). This population value is not significantly reduced from the single turn discriminability (Figure 1F, chi-squared goodness of fit, p=0.15, n=599 neurons, χ^2^=2.12, d.f.=1, odds ratio=0.79). The M2 neural population as a whole showed no laterality preference for brain hemisphere with turn encoding neurons displaying higher mean firing rates equally for ipsilateral and contralateral turns, respectively (102 contralateral preferring neurons of 203 significant turn-encoding neurons, chi-squared goodness of fit to binomial distribution, p=0.94, n=203 neurons, χ^2^=0.005, d.f.=1, odds ratio=0.99).

To assess the quality of M2 neurons’ turn discrimination, we adopted *choice probability*^64,65^ as a measure of effect size. Choice probability (CP) is simply the probability that an observer can correctly identify an outcome given a single sample from one of two distributions. It is equivalent to the area under the curve of the receiver operator characteristic^66,67^. Due to the structure of our maze and the task, sample sizes varied widely across turn locations. This measure is invariant to sample size and therefore is a particularly useful method for comparing our results.

CP confirms that M2 neurons’ encoding of turn directions is both widespread and reliable. To summarize population discrimination quality, we have adopted easily interpretable CP benchmarks of two-thirds (66%) for general discrimination and 90% for a high discrimination threshold. The second and third example neurons shown in Figure 1D-E have CP near these thresholds as examples of the average firing that leads to these levels of discrimination. On average, over half (54%) of the M2 neurons have CP exceeding 66% at any given left/right turn location, including 14% of neurons classifying at a rate exceeding 90% (Figure 1G). This is significantly higher than the CP of the same neurons with a random shuffling of turn direction identities (two-sided Kolmogorov-Smirnov test, p<0.0001 n=606 D=0.82). No neuron in the shuffled distribution discriminates above 66% (Figure 1G, max 60%). There is also no significant contextual effect elevating or suppressing turn discriminability at certain locations as all locations show statistically indistinguishable distributions of CP for left versus right turning (Figure 1H, two-sided Kruskal-Wallis test, p=0.115, χ^2^=10.24, d.f.=6).

For turn discrimination information to be accessible to downstream neurons without also passing on context, the turn choice must be encoded the same regardless of any other variable. To assess this, we pooled individual neurons’ data across all turn locations. The CP remain comparable to the location specific values, with the population counts of neurons exceeding the 66% and 90% benchmarks of discriminability only dropping from 54% to 45% and from 14% to 11% of the population, respectively (Figure 1GI).

### The Time Course of M2 Action Discrimination Involves Planning and Execution

To further delve into the time course of action discrimination we applied the CP method to each 1cm bin surrounding the first internal turn (Figure 1A, turn labeled 1). This particular ‘T’ intersection has a long approach straightaway before the turn and allows for an extended examination into left/right action discrimination. A space from 40cm before the turn apex to 15cm afterwards was analyzed.

M2 as a whole discriminates the upcoming action and continues to discriminate through action execution. M2 neuron sub-populations have peak CP values throughout the investigated epoch (Figure 2A), and a minimum of 15% of the M2 neuron population significantly differentiates the turn outcome at each time point (Figure 2B). This number increases and peaks at the turn apex with over 50% of individual M2 neurons significantly discriminating the action. The count then decreases through the end of the turn. The high reliability (>90% discrimination) neural population followed this same pattern with neuron counts ramping to the turn apex and decreasing afterwards.

**Figure 2:**
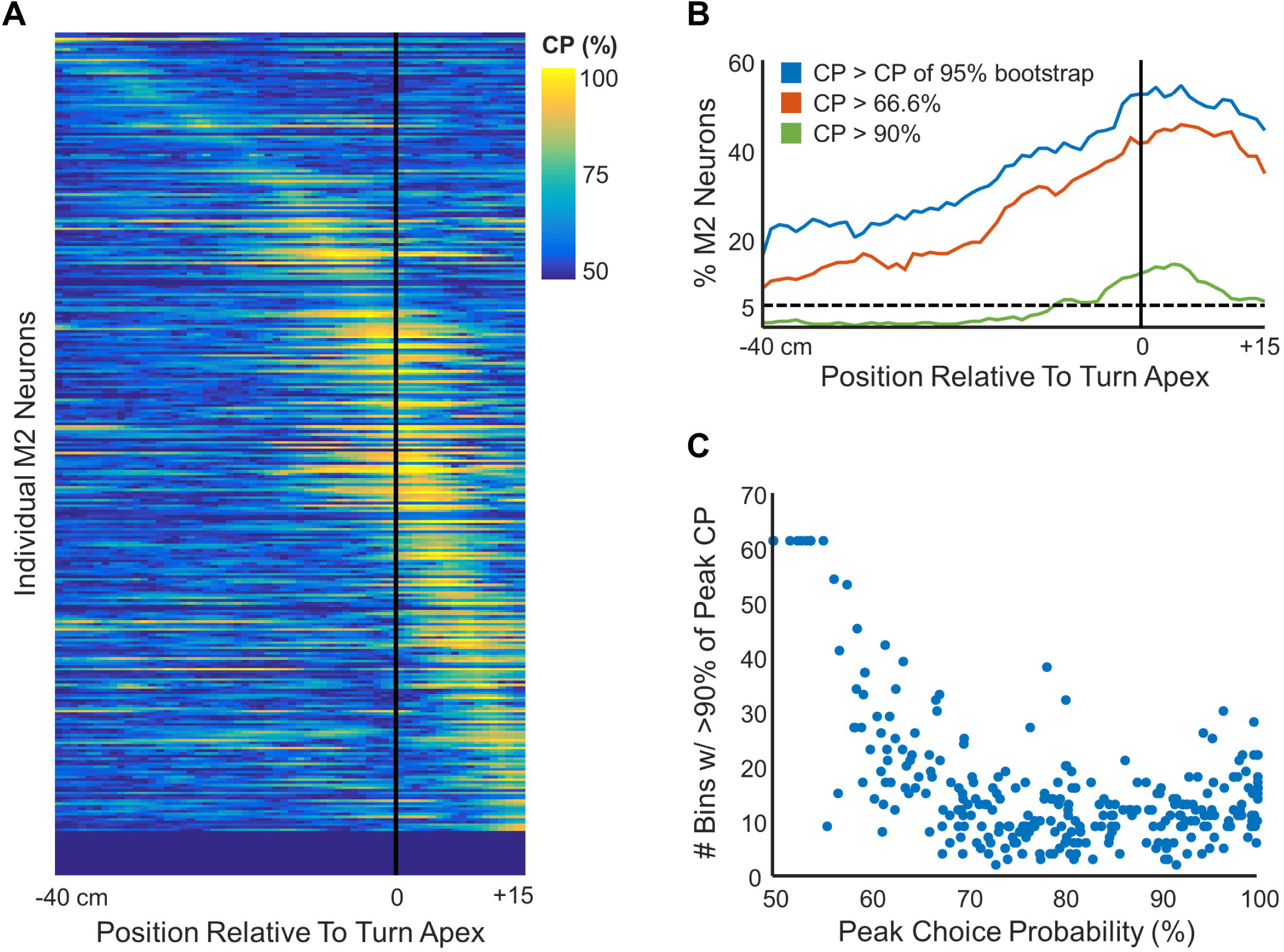
Temporal dynamics of M2 action encoding. **A)** Choice probabilities of individual neurons (rows) across space (columns) in the perievent window surrounding the first left/right turn on the internal routes (Figure 1A, labeled 1). Neurons sorted by the location of their max choice probability. Black line indicates turn apex. **B)** M2 action encoding strength as a function of time. Shown are percentages of neurons encoding the action at each spatial distance from the turn apex. Blue trace is the percentage above a 95% criteria of a bootstrapped distribution. Dotted black line is the expected chance value. Orange and green traces are percentages of neurons at the chosen thresholds of 66% and 90%, respectively. **C)** Extent of decoding period. Number of spatial bins within 10% of the maximum choice probability for each individual neuron during the time course analysis. Each dot represents one neuron.

M2 planning and action execution could either be carried out by the same neurons representing the entire period or many neurons representing the action at specific times with respect to the action. In spite of the complete temporal coverage as a population, CP values for individual neurons do not stay high for the entire period. Instead, peak CP values typically extend for a limited span of less than 20cm for any given neuron (Figure 2C) and the locations of the peak CP values vary. Given this pattern, many neurons discriminate most accurately well before the turn apex and do not significantly discriminate actions at the peak of the turn (Figure 1E middle). M2 thus contains discriminating neurons spread throughout the time sequence of planning and executing an action, and reliable discrimination through time requires an evolving ensemble of neurons.

### Widespread Representations of Spatial Context in M2

While action encoding in M2 neuron populations is clearly strong and reliable irrespective of turn location, differences between turn contexts hint that action outcome is not the sole variable represented in M2 neural activity during navigation. Considering the extensive reciprocal connectivity between M2 and both PPC and RSP, two associative cortices bearing significant spatial modulation in firing, we reasoned that M2 neurons could be integrating multiple spatial features of the environmental context in generating action-specific firing. To examine this possibility, four navigationally-relevant spatial features were tested for their impact on M2 turn-related activity – environmental/allocentric location, orientation, route being traversed, and progression within a route (1st, 2nd, or 3rd turn). Each of these variables is known to have a strong impact on the RSP, PPC or both^9, 26, 28, 33^.

To assess the impact of context (e.g., the effect of place on turn-related firing), we controlled for action (left/right turn) by calculating CP for like actions (e.g., left turns versus left turns). For some contextual variables, more than two conditions needed to be examined for discriminability. Therefore, we applied pairwise CP tests for each combination of distributions (e.g., left turn associated firing rates for each turn location against each other turn location). We present minimum, mean, and maximum CP values from these comparisons to aid in the comprehension and comparison due to the necessary differences in analyses.

Encoding of all four spatial features is widespread but with less impact on M2 activity than left versus right turning action (Figure 3A). The distributions of CP for the spatial factors in M2 are all significantly higher than chance but significantly lower than the pooled action discrimination (one-sided Kolmogorov-Smirnov test, p<0.0001, n=606, D>0.16 for all tests). The variability within individual pairwise tests can also be examined. While minimum CP for contextual variables are closer to values obtained for randomized data, minimum CP for action remain far greater than chance. Maximum CP for contextual variables are far above chance levels for the full population, but, even taking this perspective, many more neurons exhibit high CP for action discrimination. In fact, nearly no neurons reach our high discrimination threshold for any of the spatial context variables (Figure 3B). These results indicate that as a whole, M2 neuron sub-populations do accurately encode spatial and directional information, but that encoding of action (turn type) is much stronger.

**Figure 3:**
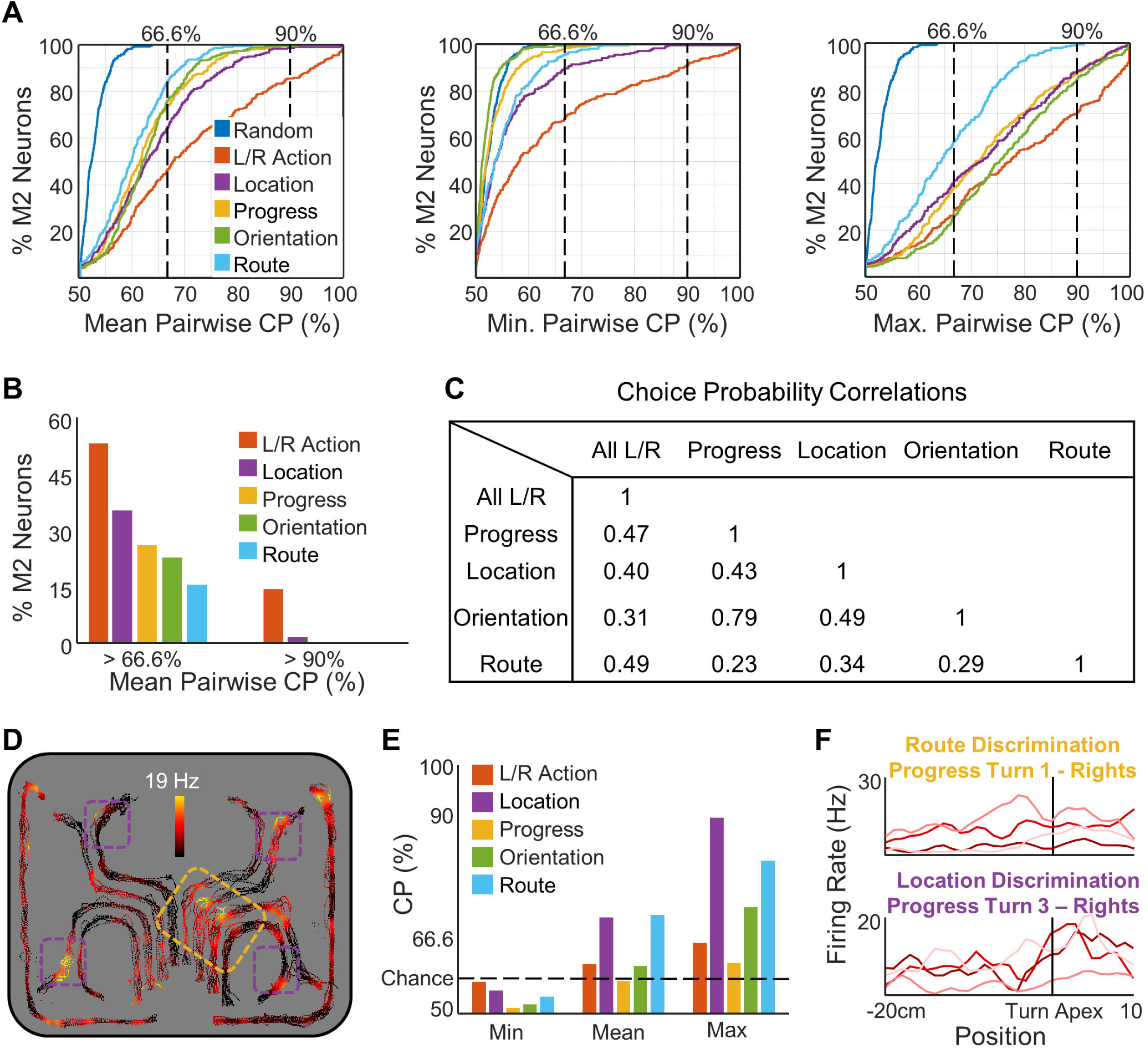
Spatial context discrimination in M2. Color labels are consistent across all of Figure 3. **A)** Cumulative density functions (CDFs) for the entire M2 populations’ CPs under different spatial conditions. Also included for comparison are CDFs of action CPs from individual locations and shuffled action CPs as in Figure 1G. Separate graphs are the mean pairwise CPs (left), minimum pairwise CPs (middle), and maximum pairwise CPs (right). **B)** Percentages of M2 neurons with mean pairwise CPs above our thresholds for each condition. **C)** Choice probability correlation table. Listed are the Pearson R correlation coefficients for the mean pairwise CPs of each pair of conditions graphed in **A. D)** Example neuron positional firing rate map color-mapped as a function of route and track position. The yellow box highlights an example of route encoding, while the purple boxes highlight location encoding. As in Figure 1B, routes are minimally translated and stretched from the actual track location to separate each map for visualization purposes. **E)** CPs from each of the three plots in A for the corresponding example neuron shown in **D. F)** Linear perievent rate maps for the highlighted examples in **D**. Examples of route discrimination from the yellow box in D (top) and location discrimination from the purple boxes in D (bottom). Only one action (right turns) shown since CPs controlled for action and would compare each action condition separately.

Given the widespread occurrence of different forms of spatial and directional information, we considered that the distribution of types of information and its co-occurrence with action discrimination may be important. Correlations of neurons’ CP across factors is shown in Figure 3C. Correlations between action encoding and each spatial factor are positive but relatively weak. The same is true for correlations between spatial factors except for the correlation between progress and orientation, which is quite high. This means that neurons that robustly encode action also tend to be neurons that encode context. Position rate maps of the routes taken are shown for an example neuron highlighting examples of conjunctive encoding in Figure 3D. Figure 3E shows the calculated mean CP for each neuron and Figure 3F highlights the mean individual firing rate vectors and separation of activity rates for different conditions. These neurons are typical examples of neurons with complex firing patterns that encode multiple factors to varying degrees.

### A Widespread but Limited Effect of Choice in M2 Action Representations

Previous work has considered a role for M2 in orienting decisions^39-44^. If the primary function of this region lies in decision-making as opposed to action planning and execution, the ability to discriminate left versus right turning action may disappear at locations where no left versus right choice exists. In our navigational task, some turns are forced (no alternative path, ‘L’) while others demand choice (‘T’). Therefore, we grouped all turn locations based on action (left or right turn) and choice context (forced vs choice). We then evaluated action discrimination at choice locations and forced-turn locations separately. There is no apparent effect on action discrimination CP due to choice context (Figure 4A). The CP distributions for forced and choice contexts did not significantly differ from each other nor from the pooled CP distribution (two-sided Kolmogorov-Smirnov test, p>0.27 n=606 D<0.09 for all tests). This result is inconsistent with lower action discrimination during forced-turn contexts and casts doubt on decision-making as the primary function of the region. Instead, the current data are consistent with M2 as a signal for upcoming and current actions.

**Figure 4:**
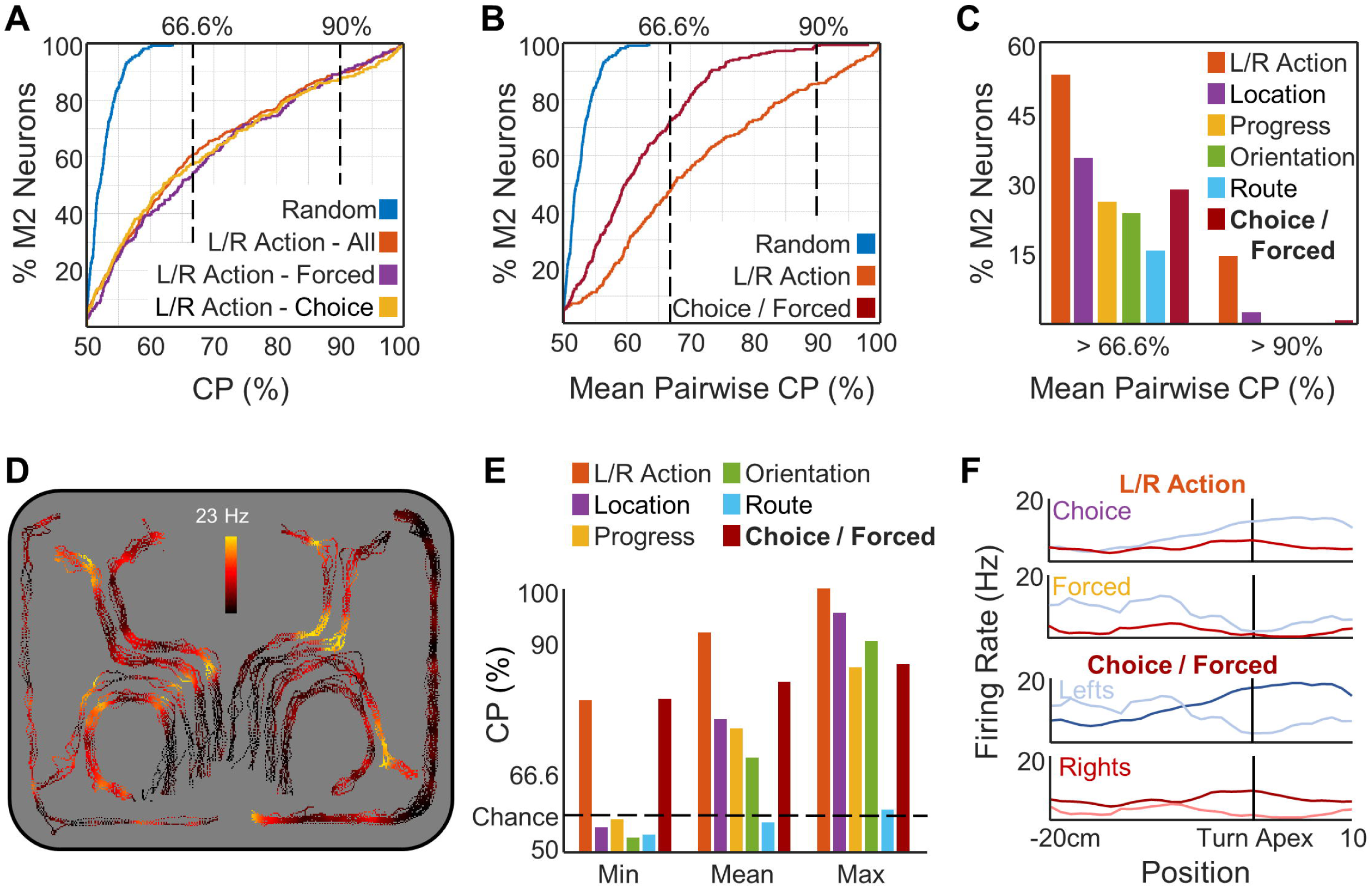
Choice context does not effect action encoding discrimination. **A)** Cumulative density functions (CDFs) for the entire M2 populations’ action CPs under choice (gold) and forced (purple) turn contexts. Also included for comparison are CDFs of action CPs from individual locations (orange) and shuffled action CPs (blue). **B)** CDF of mean pairwise CPs of choice context (maroon) with comparison CDFs of individual location action CPs (orange) and shuffled action CPs (blue). **C)** Threshold values of choice context as compared to the spatial contexts. **D)** Example neuron positional firing rate map highlighting encoding of choice context. Shown are the firing rates color-mapped as a function of route and track position. As in Figure 1B, routes are minimally translated and stretched from the actual track location to separate each map for visualization purposes. **E)** CPs from each of the three plots in Figure 3A for the example neuron shown above. **F)** Linear perievent rate maps. Perievent plots showing the mean firing rates for the action values (top) and the choice contexts (bottom) of the example neuron shown above. All comparisons for CPs were done on data being presented in the same plot.

Although action discrimination does not depend on there being a choice at all, the M2 population does discriminate between forced and choice contexts for the same action. The discriminability is similar to that of spatial factors (Figure 4BC). The position rate maps for an example action discriminating neuron with very strong modulation by choice context is highlighted in Figure 4DE. This neuron has high CP for left versus right action (higher activity for left turns) and, at the same time, fires more strongly for both left and right turns when those turns occur at locations where a left or right turn could be made (Figure 4F).

## Discussion

Prior work has considered, in detail, the firing correlates of M2 neurons to actions, action planning, and sensory integration in two-alternative, forced-choice settings^39-44^. In the present work, the task design and maze structure enabled us, for the first time, to consider these previously-identified M2 neuron correlates as they relate to the problem of navigation and integration of multiple complex spatial relationships between organisms and their environments. The ability of animals to execute fast, uninterrupted traversals of the eight internal and two external routes under these circumstances enabled us to examine a hypothesized role of M2 in the transformation of spatial cognition into action during navigation. M2 ensemble activity consistently encoded the left versus right turning action of the animal in a manner largely independent of the spatial context of any given turn site and independent of whether a choice among actions was demanded. However, action-discriminating activity itself was significantly modulated in magnitude by multiple spatial and directional variables informing turn choice at any given maze intersection. The results complement recent work outside the realm of navigation that have examined this dorsolateral prefrontal cortex sub-region and its PPC inputs as they relate to decision-making and point to the M2 as a key structure in translating more complex forms of spatial knowledge into action.

Large sub-populations of M2 neurons exhibited strong discrimination of left versus right turn actions and, in largest part, did so irrespective of spatial context given by several variables including turn location, the specific route chosen, heading direction, and route progress (i.e., turn number in a series associated with a particular route). The strength of action tuning was much higher than that for spatial and directional variables even when taking into account the larger number of possible variable states. That is, even when focusing in on the most prominent pairwise rate distinctions according to a variable such as turn location, rate distinctions for left versus right turns remain stronger across the M2 population.

By providing ample space and time between the execution of different turns and by analyzing only uninterrupted route traversals, we were also able to assess whether M2 activity dynamics are more consistent with encoding of current and planned action as opposed to a categorical decision-making process. In a decision-making process, one would expect essentially all neurons discriminating action to exhibit synchronous ramping of discriminatory activity prior to the action execution. As a population, discriminatory activity in M2 does ramp to a peak near the point of maximal angular velocity. However, locations of peak discrimination points across individual neurons varied widely in their proximity to the turn apex. For many neurons, peak discrimination of turn type not only preceded the turn apex, but also failed to discriminate turn type at the apex itself. Thus, in the context of navigation, M2 dynamics are more consistent with encoding transitions between action planning and execution rather than a deliberative process. The equal strength of predictive tuning at turn sites devoid of a decision (forced left or right) provides a second argument in favor of a relatively pure action planning and action execution role. An integrated process transitioning action plans to execution would have obvious utility in generating the fluid transitions between actions in a sequence that compose the animals’ fast, uninterrupted trajectories in the present task. Notably, disruptions in the learning of actions as part of a sequence are found subsequent to M2 lesions^68^. In this role, M2 may drive actions directly through corticospinal efferents^38^ or indirectly through intermediary structures such as primary motor cortex^37,52,69-71^. Recent data suggest that interactions between M2, primary motor cortex, ventromedial thalamus, and cerebellum may also play a key role^69-74^.

While action discrimination was found to be relatively context free, spatial and directional context significantly modulate activity rates for any single turn type. By separately examining the activity vectors surrounding all left turns and all right turns, we found that the location and directional heading of a turn relative to the full environmental space were both strong factors modulating the intensity of turn-related activity. Both the specific route taken by the animal and the position of a turn site within that route were also potent in modulating turn-related activity. The status of a turn site as ‘forced’ (only one turn type possible) versus choice (left or right turn available) proved a smaller, but not insignificant, contextual factor in modulation of M2 firing.

PPC and RSC are both major sources of afferents to the M2 sub-region of prefrontal cortex^34-36^ and have been studied extensively with respect to the forms by which they map the spatial location and directional orientation of the animal relative to the environment. PPC neurons robustly encode the location of the animal within a route space irrespective of the location of the route in the larger space of the environment^9^. Such encoding is not epiphenomenal to the tendency for a minority of PPC neurons to map specific turn types^9,27,75^. Instead, turn type specific firing for this minority population of PPC neurons is strongly modulated by the location of a turn in the environment or within a sequence^9,25,75^ such that PPC generates distinct ensemble patterns for all route locations^9,35^. While some RSC neurons generate similar spatial firing patterns for a single route placed in different regions of the larger environment, RSC ensembles generate distinct firing patterns for the same route in two different locations^28,31^. Together then, RSC and PPC together provide complementary information concerning position in a route and position in the environment. Both structures also contain small populations of head direction cells^26,28,76,77^ and discrimination of two routes taken through the same location has been reported for both PPC and RSC ensembles^33,78^. Thus, PPC and RSC stand as two likely sources of modulation of M2 neurons according to the environmental location of a turn, the environment-referenced set of head orientations of the animal as it proceeds through a turn, the shape of the full route taken by the animal, and the ordering of turns within a route.

Our use of CP analysis to characterize tuning of M2 neurons to left/right turning actions and to spatial and orientational variables leads us to conclude that both actions and the context in which they occur are encoded in M2 populations. This raises the question of whether M2 functions to encode the context of actions, encodes only upcoming and current actions, or both. Here, distinct differences in the strength of encoding of different variables perhaps offers an answer. Under essentially all circumstances, modulation of single neuron and ensemble activity rates distinguishing actions is clearly greater than that for the spatial and choice variables considered. Given this circumstance, a downstream target that can directly mediate actions could, through degeneration, produce the same activity patterns related to left or right turning in response to largely overlapping input patterns for different spatial contexts. In this way, a pure action signal could be derived despite the variability in patterning of M2 neurons according to spatial context. A second way to accomplish the same outcome, one that cannot be addressed here, is through specificity of projections. M2 neurons that strongly discriminate left/right turning actions without discriminating context could be biased in their efferent connectivity. An argument against this model is that tuning for action and spatial context in M2 is positively correlated. Nevertheless, recent work^79^ does show that M2 populations segregate in lateralization of their action tuning according to whether they project intracortically or to brainstem targets. Projection specificity of M2 neurons with context-independent encoding of action will be a question of importance in future work aimed at determining how M2 signals are interpreted by its efferent targets.

M2 efferents reach many brain regions and so one feature of M2 function could be to supply a conjunctive efference-copy that reflects the full context of actions and the spatial and choice context in which they occur. Action tuning observed in RSC and PPC, for example, could reflect the efferents of M2 neurons to these regions. We conclude, however, that M2’s role is best understood as a part of a sensory and spatial associative cortex network that outputs preparatory motor actions. This view aligns closely with that posited in prior work emphasizing experiments wherein spatial context was not considered and emphasizes M2’s situated place in the PPC and RSP cortical network. In this respect, it follows that M2 tuning to spatial context may be a remnant of an integration process in which spatial information is transformed into an appropriate motor plan.

We conclude that M2 ensemble activity discriminates, in a manner largely independent of spatial context, the specific motor acts associated with navigation constrained to paths. We also conclude that such activity can contribute to both the planning and execution of action. The implication of M2 in contributing to action planning in navigation is consistent with prior work emphasizing the role of this particular sub-region of prefrontal cortex in generating activity patterns predictive of upcoming choices during delay periods preceding a single choice point^41^. It is also consistent with work demonstrating that M2 is required for action planning^73^ cued by single modality sensory cues of multiple types^80^. The present work demonstrates that such activity is distributed across a broad expanse of space and time prior to peak turning behavior and can therefore manifest as a continuous process during active and uninterrupted navigation through a series of left/right turn choices. This distinguishes the present finding from prior work examining M2 activity during delay periods absent locomotion^41^ and, furthermore, dissociates action type from reward location as the relevant variable predicted by M2 during delay periods of navigation tasks. Finally, the ramping of discriminatory activity to a peak at the height of turns is suggestive of a network process internal to M2 that continually mediates transitions from planned to executed actions.

## Materials and Methods

### Subjects

Subjects were 7 adult male Sprague-Dawley rats. From these rats, a total of 303 neurons were recorded (73 left, 230 right; for per animal counts, see Supplemental Table 1). Rats were housed individually and kept on a 12-h light/dark cycle. Prior to experimentation, animals were habituated to the colony room and handled for 1–2 weeks. During training and experimentation, rats were food-restricted with weights maintained at 85–95% of their free-feeding weight. Water was available continuously. Rats were required to reach a minimum weight of 350g (5–10 months of age) before surgery and subsequent experimentation. All experimental protocols adhered to AALAC guidelines and were approved by the IACUCs through either the UCSD Animal Care Program or the Scripps Research Institute.

### Apparatus

Behavioral tasks were conducted using a ‘triple-T’ track maze. The track (Figure 1A, left panel; 8-cm-wide pathways, overall 1.6m × 1.25m in length and width, painted black) stood 20cm high in the middle of a large recording room. The track edges were only 2cm in height, allowing an unobstructed view of the environment’s boundaries and associated distal visual cues.

### Behavior

Rats were habituated to the ‘triple-T’ maze during two 30-minute periods of free exploration. Animals were then trained to run in an uninterrupted fashion from the start location to one of eight potential goal sites distributed near the perimeter of the full maze (Figure 1A, yellow lines). These eight ‘internal’ routes consisted of straight sections interleaved with three left or right turns prior to a full stop at a goal location. The animal must then travel along the perimeter of the maze to the start location to begin a new trial. The ‘perimeter’ routes selected were typically those yielding the shortest distance to the start site. Internal route lengths were 140cm in total length, with turns at 51cm, 87cm and 118cm. Perimeter routes varied considerably in length according to the distances between goal locations and the start site. If warranted, reward (a half piece of Cheerios cereal) was made available at the goal sites. Over 1–2 weeks, animals were trained by approximation to make route traversals between goal sites. Over at least two additional weeks, animals were trained by simple trial and error to a criterion of 80% ballistic (uninterrupted) route traversals. Animals were surgically implanted only after this level of task performance had been achieved.

### Reward Schedules

Multiple reward schedules were used across the set of animals. For the visit-all-8 reward paradigm, the animal was rewarded at all eight goal locations, but needed to visit all locations before rewards were reset at all reward locations. For the visit-all-4 reward paradigm, only the far four routes and reward locations were included. For these animals, the other four routes were blocked and the animals never had access to those goal sites. In the high-low reward paradigm, two locations out of the eight were randomly chosen to be rewarded for each recording. One location contained one half Cheerio reward and the other one quarter. After 20 minutes, two more randomly selected locations were chosen to be rewarded in the same high-low fashion and the recording continued for 20 more minutes. Under the high-low paradigm, rats primarily visited the high-reward locations, effectively limiting sampling to three internal routes.

### Surgery

Rats were surgically implanted with stereotrode or tetrode arrays (twisted sets of two 25µm tungsten wires or four 12.5µm nichrome wires) inserted into custom-built microdrives (four to eight arrays per microdrive). Rats were implanted unilaterally or bilaterally with one microdrive per hemisphere into M2. Rats were anesthetized with isoflurane and positioned in a stereotaxic device (Kopf Instruments). Following craniotomy and resection of dura mater, microdrives were implanted relative to bregma, centered at (A/P 2.5mm, M/L ±1.2mm, D/V −0.5mm).

### Recordings

After a one-week recovery from surgery, animals were retrained for at least one week before beginning recordings. This was to ensure adequate behavior and running ability with the new weight of the implant. All recordings were from animals that were well trained on the task. Electrodes were moved ventrally in 40-80 µm increments between recordings to maximize the number of distinct units collected. Omnetics connectors were connected to the microdrives for all animals. One of two recording systems was used for data collection. In one, utilized for five animals, each Omnetics connector was connected to a single amplifying headstage (1X gain, NB Labs). A tether led to a bank of amplifiers (Lynx-8, Neurolynx) and then fed into an acquisition computer running the AD system (courtesy of Loren Frank and Matt Wilson, MIT). Signals were filtered at 0.6 to 6kHz, amplified between 1,000 to 20,000X, and digitized at 32kHz. For the other two animals, each microdrive had two electrical interface boards connected to a single amplifying headstage (20X, Triangle Biosystems). A tether led to a set of preamplifiers (50X) and a high pass filter (>150Hz). Signals then fed into the acquisition computer running Plexon SortClient software and were filtered at 0.45–9kHz, further amplified 1–15X, and digitized at 40kHz. Total amplification regardless of system was a total of 1,000–20,000X). Single unit action potential waveforms were isolated in either XClust (courtesy of Loren Frank and Matt Wilson, MIT) or Plexon OfflineSorter software. Waveform parameters used were peak height, peak valley, energy, and principal components. Waveform clusters appearing to overlap with the amplitude threshold set for collection were discarded to avoid collection of neurons with partial spiking data. Waveform amplitudes were monitored to ensure systematic fluctuation over time did not produce confounds in isolating single units. Recordings lasted approximately 30-60 minutes. No neurons were excluded from analysis, even if activity was minimal.

### Position Tracking

Animal position was tracked using a camera attached to the ceiling above the recording room floor. Either Dragon Tracker hardware (AD recording system) or Plexon’s CinePlex Studio software was used to detect LED lights fitted to the animal’s surgical implant before each recording. Light positions were captured at 60Hz. Accurately identifying and defining the actions of a freely navigating animal for analysis can be difficult due to the wide variation of movements. To control for such variation in analyzing action potential firing rates, we filtered the animal tracking data for smooth, uninterrupted traversals of entire routes. This procedure permitted us to examine action potential firing rate data associated with stereotypical movements (forward running, turning) through all sections of all routes. The defined routes were the eight internal routes toward potential reward sites or perimeter routes leading to the internal entrance (Figure 1B). As perimeter routes from the four goal locations nearest the start point were excessively short, the perimeter routes used for analysis were the two two-turn routes returning the animal from the first four internal route end-points to the start point. Each clean run was linearized by mapping tracking locations to the nearest 1cm bin of a template of the traversed routes.

### Histology

Animals were perfused with 4% paraformaldehyde under deep anesthesia. Brains were removed and sliced into 50µm sections and Nissl-stained to reveal the final depth of electrode wires in M2. Microdrive depth monitored across recordings and final electrode depth as observed in histology were compatible in all animals reported and a schematic of fiber tracks is shown in Figure 1C.

### Identification of Fluid, Efficient Route Traversals

To identify uninterrupted runs for individual routes, a multistep process using custom MATLAB graphical user interfaces was used. First, a user defines starting and ending gates for each route. Then the program finds all runs crossing these locations with sustained running speeds of 3cm/s or greater throughout. Finally, a researcher uses the interface to verify that all such identified runs did not contain either obvious interruptions in locomotion or significant alterations from the stereotyped path seen across multiple runs. The results of this process can be seen in Figure 1B. By this method, stalled track traversals, reward periods, and other position data captured between runs are not conflated with data with controlled action and spatial values.

### Positional Firing Rate Maps

To characterize the firing activity of the M2 neurons, we calculated individual neurons’ positional firing rates by dividing the total number of spikes of each neuron at each location by the total occupancy time at each location. Only position samples included in identified ballistic route traversals were used in such calculations and rate vectors for each route were generated separately. Positional firing maps were smoothed using a 2D convolution with a Gaussian filter with a standard deviation of 2cm that also accounts for bins with no occupancy^81^.

For each recording, custom MATLAB software is utilized to generate a spatial template matching the average trajectory of the animal along each route in the horizontal (2D) plane. This approach ensures the best possible matching of animal behavior and positions taken across recordings and trials. The beginnings and ends of each straight run and turn peaks are marked and equally sized bins are interpolated between marked locations. After establishing the template, each tracking position sample is mapped to the nearest template bin. Firing rates for each bin of each route are calculated by summing the number of spikes in each template bin and dividing by the total amount of time the bin was occupied. Mean firing rates are calculated by summing mean rates at each location and dividing by the number of runs.

### Peri-event Mean Firing Rates

Peri-event mean firing rates were used to calculate choice probabilities at turns. For the defined linear space surrounding positions peak turning behavior occurs, firing rates for all occupied bins of linearized positional rate vectors were averaged for each trial. Bin sizes were 1cm and the length of the pre-turn and post-turn spaces were 20cm and 10cm respectively. This approach allowed us to make direct comparisons of activity rates for all pre-turn and post-turn spaces across turn types (left versus right) and for the same turn type at different maze locations.

### Choice Probability

Choice probability (CP) is a metric coined by Britten et al.^64^ It is the probability that an observer can correctly identify an outcome given a single sample from one of two distributions. It is equivalent to the area under the curve of the receiver operator characteristic^66-67^. It can be calculated from the U statistic of the Mann Whitney U test of two distributions,

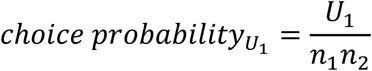

where *n*_*1*_ and *n*_*2*_ are the number of samples in distributions 1 and 2, respectively. The difference of the value of this metric from chance (50%) is symmetric but depends on the order of the magnitudes of the medians of the two distributions. For our purposes, the higher magnitude of the distributions was not important, only the separation. Because of this, the maximum choice probability,

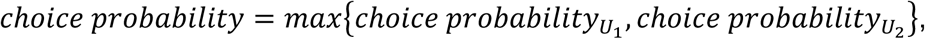

was always selected.

For all peri-event action choice probabilities (based on spike rates for left versus right turns), peri-event mean firing rates were calculated from the distributions of routes with left and right actions. For the time course analysis, choice probabilities were computed from firing rates of a particular bin instead of mean values across bins. To control for the effect of noisier data at this fine granularity and multiple comparisons, the significance level for the time course test was established by a bootstrapping procedure. We shuffled the left/right action identities of the same runs analyzed for each bin and neuron to create 1000 shuffled left/right datasets for each bin and for each neuron. Choice probabilities from this shuffled distribution were calculated and used to establish a p < 0.05% criterion for choice probability values.

For contextual choice probabilities (e.g., the effect of place on turn-related firing), we always controlled for action (left/right turn) by comparing only like actions (e.g., left turns versus left turns). For many contextual factors, more than two conditions needed to be examined for discriminability. To preserve the ease of interpretation, we decided upon pairwise choice probability calculations for each combination of distributions. For progress within a route, this consisted of 12 comparisons: 1st, 2nd, and 3rd turns of the 8 internal routes. For location, we controlled for route progress and therefore had up to 26 comparisons: between four 3rd turn locations (6 combinations), and between two 2nd turn locations (1 combination) for each of left and right turns. For route, we controlled for location and progress by analyzing only the 1st turn common to all 8 internal paths. We therefore had 12 comparisons: 6 from four left routes and 6 from 4 right routes. For orientation we had 12 comparisons as there were four possible trajectories for lefts and rights (north-going to west-going, north to east, south to west, south to east). Finally, for choice, we only had two comparisons, forced versus choice, for lefts and rights that occurred at locations where the animal could or could not execute more than one type of turn. We present minimum, mean, and maximum values from these comparisons to aid in the comprehension and comparison due to the necessary differences in analyses.

### Statistical tests

Nonparametric tests were used throughout to avoid assumptions of normality in the data. The Mann Whitney U test was used to evaluate the statistical significance of peri-event mean firing rates between left and right actions. Fisher’s exact test was used to assess the quantities of significant neurons for the first turn alone (common to all 8 internal paths) and for all turns pooled (all internal and perimeter turns). A Chi-Squared Goodness of Fit test against an unbiased binomial distribution was used to assess deviation from chance (50%) for laterality preference of M2 neurons. The Kruskal-Wallis test was utilized to evaluate whether action choice probabilities from separate locations were from different distributions. The Kolmogorov-Smirnov test was used to assess if pairwise distributions were significantly different for CP distributions. No statistical methods were used to predetermine sample sizes. However, based on similar sample sizes reported in previous publications, we believe we have adequate power (0.8) or greater to detect significant effects.

## Data and code availability

The data that support the findings of this study and code used for analysis are available from the corresponding author upon reasonable request.

## Acknowledgements

This work was supported by the National Science Foundation (IOS-1149718).

The authors wish to thank the following people for technical assistance, discussion of the data, and editing of the original manuscript: Kara Papaefthimiou, Ari Kappel, Belinda La, Cody Walters, Emily Pham, Jinho Chung, Natalie Tongprasearth, Emily Tao, Scott Ragland, Eran Mukamel, Laleh Quinn, Lara Rangel, Andrea Chiba, Andrew Alexander, Laura Shelley, and Alex Johnson.

The authors declare no competing financial interest.

---

**Supplemental Table 1:**
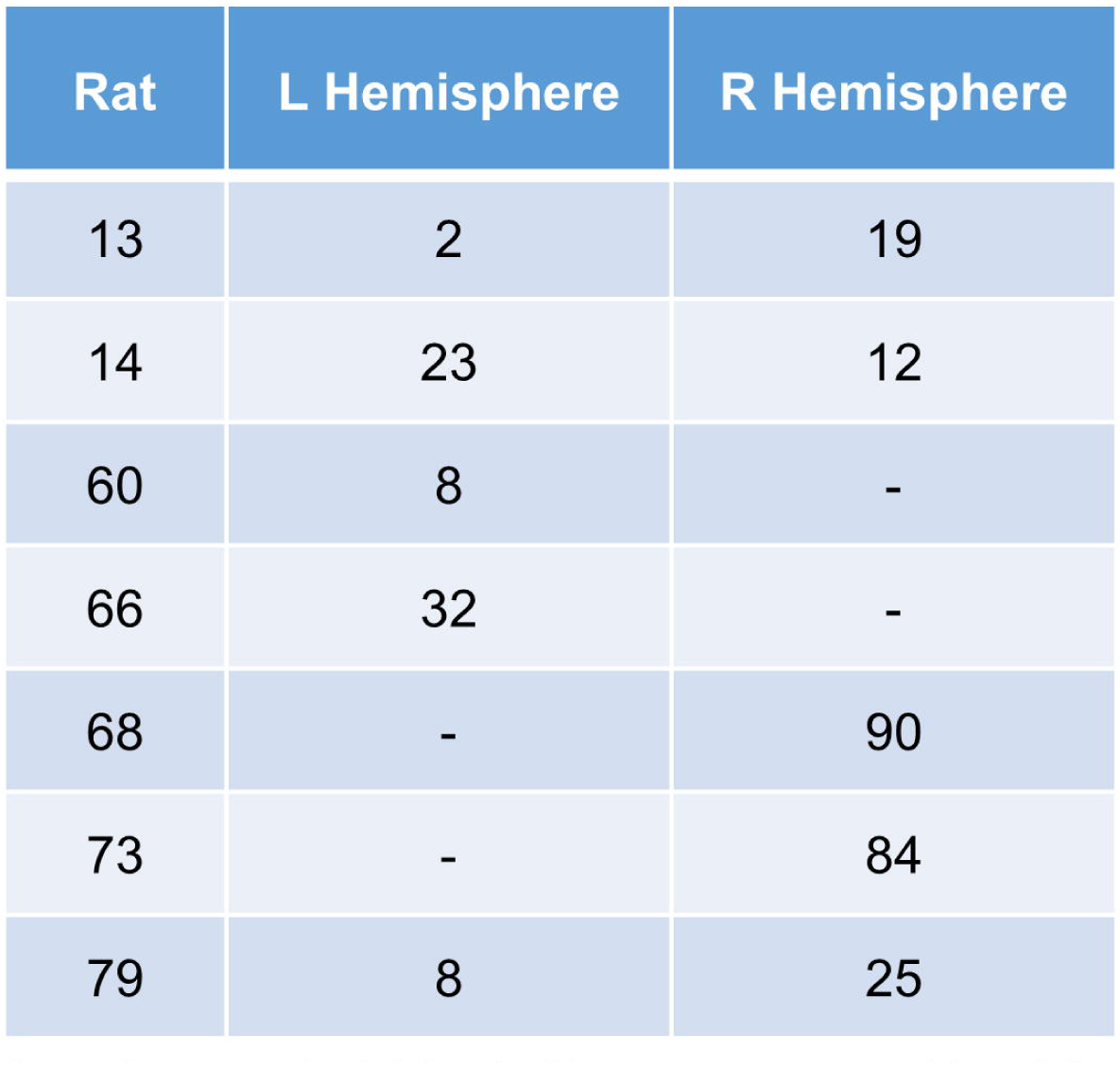
Neuron counts. Listed in each row is the identifier index and neuron coun ts by hemisphere for each rat in the study.

**Supplemental Figure 1:**
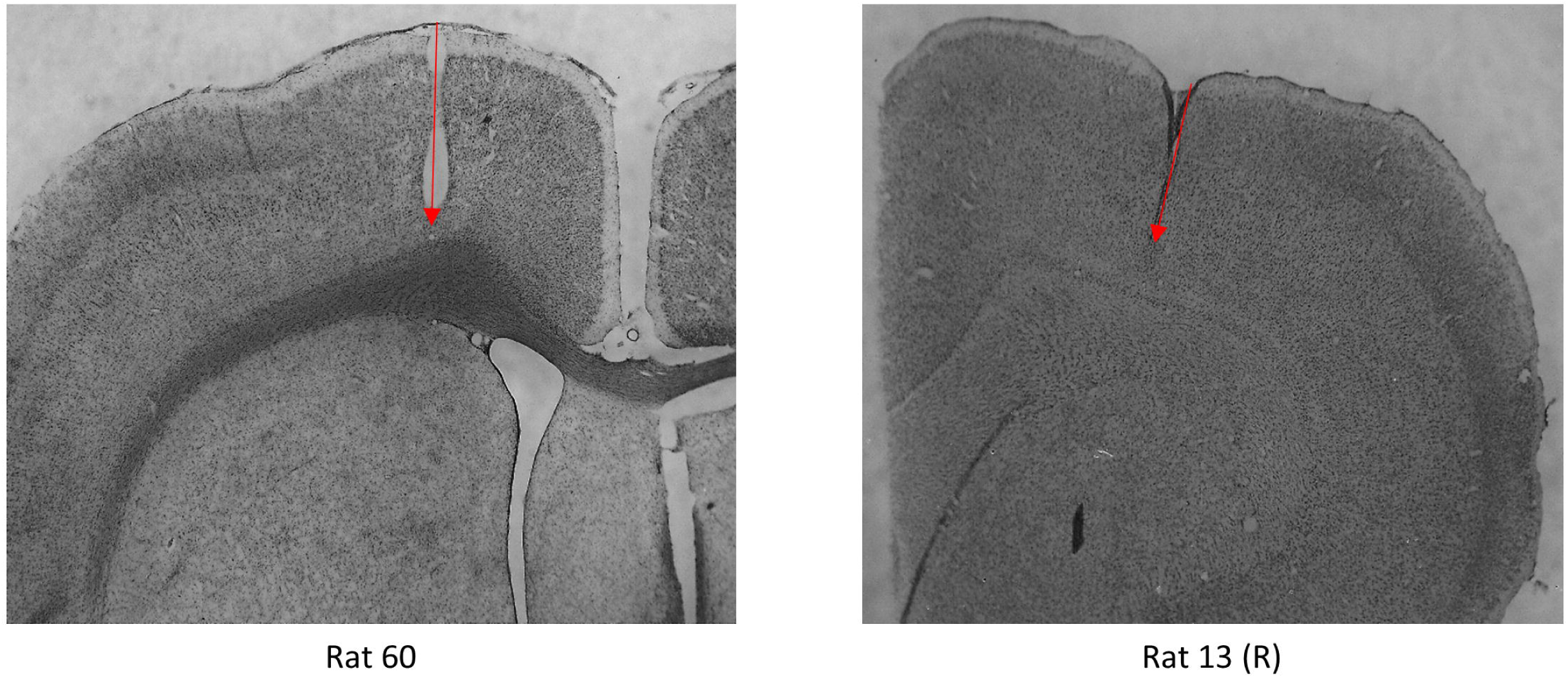
Histology. Recordings of M2 neurons (N = 303) were obtained from nine bundles n animals. Numbers of total recorded neurons are included in Supplemental Table 1. Example y photos. Red arrows depict tracks left by the bundles and their approximate endpoints.

**Supplemental Figure 2:**
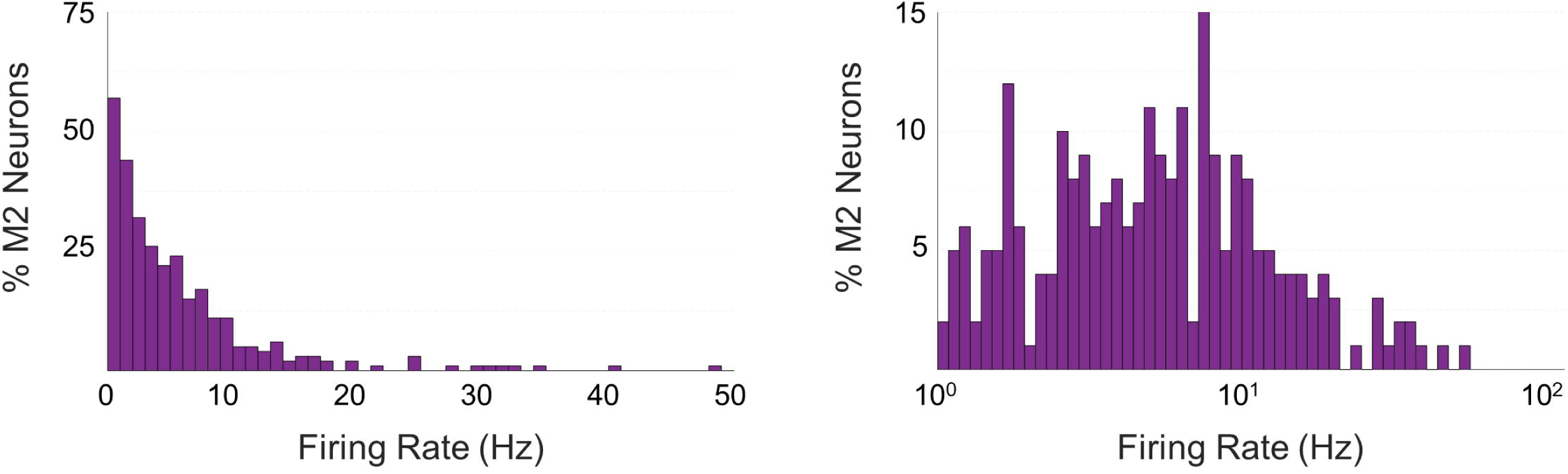
Firing rate distributions. Mean firing rate histograms for the 303 M2 neurons in both linear (left) and log-linear scales (right).

## Notes

#### Summary of Updates

Stats updated.

## References

1. Bassett JP, Taube JS. (2001) Neural correlates for angular head velocity in the rat dorsal tegmental nucleus. J Neurosci. 21(15):5740–51.

2. Wang C, Chen X, Lee H, Deshmukh SS, Yoganarasimha D, Savelli F, Knierim JJ (2018) Egocentric coding of external items in the lateral entorhinal cortex. Science 23;362(6417):945–949.

3. LaChance PA, Todd TP, Taube JS (2019) A sense of space in postrhinal cortex. Science 365(6449).

4. Dale A, Cullen KE (2019) The Ventral Posterior Lateral Thalamus Preferentially Encodes Externally Applied Versus Active Movement: Implications for Self-Motion Perception. Cereb Cortex 29(1):305–318

5. Hinman JR, Chapman GW, Hasselmo ME (2019) Neuronal representation of environmental boundaries in egocentric coordinates. Nat Commun. 10(1):2772

6. O’Keefe J, Dostrovsky J. (1971) The hippocampus as a spatial map. Preliminary evidence from unit activity in the freely-moving rat. Brain Res. 34(1):171–5.

7. Taube JS, Muller RU, Ranck JB Jr. (1990) Head-direction cells recorded from the postsubiculum in freely moving rats. I. Description and quantitative analysis. J Neurosci.10(2):420–35.

8. Hafting T, Fyhn M, Molden S, Moser MB, Moser EI (2005) Microstructure of a spatial map in the entorhinal cortex. Nature 436(7052):801–6.

9. Nitz DA. (2006) Tracking route progression in the posterior parietal cortex. Neuron. 2006 49(5):747–56.

10. Savelli F, Yoganarasimha D, Knierim JJ. (2008) Influence of boundary removal on the spatial representations of the medial entorhinal cortex. Hippocampus. 18(12):1270–82

11. Stewart S, Jeewajee A, Wills TJ, Burgess N, Lever C. (2013) Boundary coding in the rat subiculum. Philos Trans R Soc Lond B Biol Sci. 369(1635).

12. Olson JM, Tongprasearth K, Nitz DA (2017) Subiculum neurons map the current axis of travel. Nat Neurosci. 2017 20(2):170–172.

13. Bos JJ, Vinck M, van Mourik-Donga LA, Jackson JC, Witter MP, Pennartz CMA (2017) Perirhinal firing patterns are sustained across large spatial segments of the task environment. Nat Commun. 8:15602

14. Peyrache A, Schieferstein N, Buzsáki G (2017) Transformation of the head-direction signal into a spatial code. Nat Commun. 8(1):1752

15. Høydal ØA, Skytøen ER, Andersson SO, Moser MB, Moser EI (2019) Object-vector coding in the medial entorhinal cortex. Nature. 568(7752):400–404.

16. Vogt BA, Miller MW (1983) Cortical connections between rat cingulate cortex and visual, motor, and postsubicular cortices. J Comp Neurol. 216(2):192–210.

17. Van Groen T, Wyss JM (1990) Connections of the retrosplenial granular a cortex in the rat. J Comp Neurol. 300(4):593–606.

18. Burwell RD, Amaral DG (1998) Cortical afferents of the perirhinal, postrhinal, and entorhinal cortices of the rat. J Comp Neurol. 398(2):179–205.

19. Van Groen T, Wyss JM (2003) Connections of the retrosplenial granular b cortex in the rat. J Comp Neurol. 463(3):249–63.

20. Agster KL, Burwell RD (2009) Cortical efferents of the perirhinal, postrhinal, and entorhinal cortices of the rat. Hippocampus. 19(12):1159–86.

21. Czajkowski R, Sugar J, Zhang SJ, Couey JJ, Ye J, Witter MP (2013) Superficially projecting principal neurons in layer V of medial entorhinal cortex in the rat receive excitatory retrosplenial input. J Neurosci. 33(40):15779–92.

22. Olsen GM, Witter MP (2016) Posterior parietal cortex of the rat: Architectural delineation and thalamic differentiation. J Comp Neurol. 524(18):3774–3809.

23. Olsen GM, Ohara S, Iijima T, Witter MP (2017) Parahippocampal and retrosplenial connections of rat posterior parietal cortex. Hippocampus. 27(4):335–358.

24. Yamawaki N, Corcoran KA, Guedea AL, Shepherd GMG, Radulovic J (2019) Differential Contributions of Glutamatergic Hippocampal→Retrosplenial Cortical Projections to the Formation and Persistence of Context Memories. Cereb Cortex. 29(6):2728–2736.

25. McNaughton BL, Mizumori SJ, Barnes CA, Leonard BJ, Marquis M, Green EJ. (1994) Cortical representation of motion during unrestrained spatial navigation in the rat. Cereb Cortex. 4(1):27–39.

26. Cho J, Sharp PE (2001) Head direction, place, and movement correlates for cells in the rat retrosplenial cortex. Behav. Neurosci. 115(1):3–25.

27. Whitlock JR, Pfuhl G, Dagslott N, Moser MB, Moser EI (2012) Functional split between parietal and entorhinal cortices in the rat. Neuron. 73(4):789–802.

28. Alexander AS, Nitz DA (2015) Retrosplenial cortex maps the conjunction of internal and external spaces. Nat Neurosci. 18(8):1143–51.

29. Alexander AS, Nitz DA (2017) Spatially Periodic Activation Patterns of Retrosplenial Cortex Encode Route Sub-spaces and Distance Traveled. Curr Biol. 27(11):1551–1560.

30. Mimica B, Dunn BA, Tombaz T, Bojja VPTNCS, Whitlock JR (2018) Efficient cortical coding of 3D posture in freely behaving rats. Science. 362(6414):584–589.

31. Wilber AA, Clark BJ, Forster TC, Tatsuno M, McNaughton BL. (2014) Interaction of egocentric and world-centered reference frames in the rat posterior parietal cortex. J Neurosci. 34(16):5431–46.

32. Yang FC, Jacobson TK, Burwell RD (2017) Single neuron activity and theta modulation in the posterior parietal cortex in a visuospatial attention task. Hippocampus. 27(3):263–273.

33. Miller AMP, Mau W, Smith DM (2019) Retrosplenial Cortical Representations of Space and Future Goal Locations Develop with Learning. Curr Biol. 29(12):2083–2090.

34. Reep RL, Chandler HC, King V, Corwin JV (1994) Rat posterior parietal cortex: topography of corticocortical and thalamic connections. Exp Brain Res. 100(1):67–84.

35. Nitz D (2009) Parietal cortex, navigation, and the construction of arbitrary reference frames for spatial information. Neurobiol Learn Mem. 2009 Feb;91(2):179–85.

36. Yamawaki N, Radulovic J, Shepherd GM (2016) A Corticocortical Circuit Directly Links Retrosplenial Cortex to M2 in the Mouse. J Neurosci. 36(36):9365–74.

37. Gu X, Staines WA, Fortier WP (1999) Quantitative analyses of neurons projecting to primary motor cortex zones controlling limb movements in the rat. Brain Res. 835:175–187.

38. Ueno M, Nakamura Y, Li J, Gu Z, Niehaus J, Maezawa M, Crone SA, Goulding M, Baccei ML, Yoshida Y (2018) Corticospinal Circuits from the Sensory and Motor Cortices Differentially Regulate Skilled Movements through Distinct Spinal Interneurons. Cell Rep. 23(5):1286–1300.

39. Barthas F, Kwan AC (2017) Secondary Motor Cortex: Where ‘Sensory’ Meets ‘Motor’ in the Rodent Frontal Cortex. Trends Neurosci. 40(3):181–193.

40. Erich JC, Bialek M, Brody CD (2011) A cortical substrate for memory-guided orienting in the rat. Neuron. 72(2):330–43.

41. Sul JH, Jo S, Lee D, Jung MW (2011) Role of rodent secondary motor cortex in value-based action selection. Nat Neurosci. 14(9):1202–8.

42. Murakami M, Vicente MI, Costa GM, Mainen ZF (2014) Neural antecedents of self-initiated actions in secondary motor cortex. Nat Neurosci. 17(11):1574–82.

43. Li N, Daie K, Svoboda K, Druckmann S (2015) Robust neuronal dynamics in premotor cortex during motor planning. Nature. 532(7600):459–64.

44. Hanks TD, Kopec CD, Brunton BW, Duan CA, Erlich JC, Brody CD (2015) Distinct relationships of parietal and prefrontal cortices to evidence accumulation. Nature. 520(7546):220–3.

45. Peters AJ, Chen SX, Komiyama T (2014) Emergence of reproducible spatiotemporal activity during motor learning. Nature. 510(7504):263–7.

46. Siniscalchi MJ, Phoumthipphavong V, Ali F, Lozano M, Kwan AC (2016) Fast and slow transitions in frontal ensemble activity during flexible sensorimotor behavior. Nat Neurosci. 19(9):1234–42.

47. Chen TW, Li N, Daie K, Svoboda K (2017) A Map of Anticipatory Activity in Mouse Motor Cortex. Neuron. 94(4):866–879.

48. Jung MW, Qin Y, McNaughton BL, Barnes CA (1998) Firing characteristics of deep layer neurons in prefrontal cortex in rats performing spatial working memory tasks. Cereb Cortex. 8(5):437–50.

49. Pratt WE, Mizumori SJ (2001) Neurons in rat medial prefrontal cortex show anticipatory rate changes to predictable differential rewards in a spatial memory task. Behav Brain Res. 123(2):165–83

50. Jones MW, Wilson MA (2005) Theta rhythms coordinate hippocampal-prefrontal interactions in a spatial memory task. PLoS Biol. 3(12):e402.

51. Euston DR, McNaughton BL (2006) Apparent encoding of sequential context in rat medial prefrontal cortex is accounted for by behavioral variability. J Neurosci. 26(51):13143–55.

52. Narayanan NS, Laubach M (2006) Top-down control of motor cortex ensembles by dorsomedial prefrontal cortex. Neuron. 2006 Dec 7;52(5):921–31.

53. Kargo WJ, Szatmary B, Nitz DA (2007) Adaptation of prefrontal cortical firing patterns and their fidelity to changes in action-reward contingencies. J Neurosci. 2007 Mar 28;27(13):3548–59.

54. Sul JH, Kim H, Huh N, Lee D, Jung MW (2010) Distinct roles of rodent orbitofrontal and medial prefrontal cortex in decision making. Neuron. 66(3):449–60.

55. Durstewitz D, Vittoz NM, Floresco SB, Seamans JK (2010) Abrupt transitions between prefrontal neural ensemble states accompany behavioral transitions during rule learning. Neuron. 2010 May 13;66(3):438–48.

56. Cowen SL, Davis GA, Nitz DA (2012) Anterior cingulate neurons in the rat map anticipated effort and reward to their associated action sequences. J Neurophysiol. 2012 May;107(9):2393–407.

57. Horst NK, Laubach M (2012) Working with memory: evidence for a role for the medial prefrontal cortex in performance monitoring during spatial delayed alternation. J Neurophysiol. 108(12):3276–88

58. Hyman JM, Whitman J, Emberly E, Woodward TS, Seamans JK (2013) Action and outcome activity state patterns in the anterior cingulate cortex. Cereb Cortex 23(6):1257–68.

59. Ito HT, Zhang SJ, Witter MP, Moser EI, Moser MB (2015) A prefrontal-thalamo-hippocampal circuit for goal-directed spatial navigation. Nature. 522(7554):50–5.

60. Powell NJ, Redish AD (2016) Representational changes of latent strategies in rat medial prefrontal cortex precede changes in behaviour. Nat Commun. 7:12830.

61. Malagon -Vina H, Ciocchi S, Passecker J, Dorffner G, Klausberger T (2018) Fluid network dynamics in the prefrontal cortex during multiple strategy switching. Nat Commun. 9(1):309

62. Yu JY, Liu DF, Loback A, Grossrubatscher I, Frank LM (2018) Specific hippocampal representations are linked to generalized cortical representations in memory. Nat Commun. 2018 Jun 7;9(1):2209.

63. Buzsaki G, Mizuseki K (2014) The log-dynamic brain: how skewed distributions affect network operations. Nat. Rev. Neurosci. 15:264–78.

64. Britten KH, Newsome WT, Shadlen MN, Celebrini S, Movshon JA (1996) A relationship between behavioral choice and the visual responses of neurons in macaque MT. Vis. Neurosci. 13:87–100.

65. Erlich JC, Bialek M, Brody CD (2011) A cortical substrate for memory-guided orienting in the rat. Neuron (72)330–43.

66. Green DW, Swets JA (1966) Signal detection theory and psychophysics. Wiley Publishing.

67. Bamber D (1975) The area above the ordinal dominance graph and the area below the receiver operating characteristic graph. J. Math. Psychol. 12:387–415.

68. Ostlund SB, Winterbauer NE, Balleine BW (2009) Evidence of action sequence chunking in goal-directed instrumental conditioning and its dependence on the dorsomedial prefrontal cortex. J Neurosci. 29(25):8280–7.

69. Smith NJ, Horst NK, Liu B, Caetano MS, Laubach M (2010) Reversible Inactivation of Rat Premotor Cortex Impairs Temporal Preparation, but not Inhibitory Control, During Simple Reaction-Time Performance. Front Integr Neurosci. 4:124

70. Economo MN, Viswanathan S, Tasic B, Bas E, Winnubst J, Menon V, Graybuck LT, Nguyen TN, Smith KA, Yao Z, Wang L, Gerfen CR, Chandrashekar J, Zeng H, Looger LL, Svoboda K (2018) Distinct descending motor cortex pathways and their roles in movement. Nature. Nov;563(7729):79–84.

71. Svoboda K, Li N (2018) Neural mechanisms of movement planning: motor cortex and beyond. Curr Opin Neurobiol. 49:33–41.

72. Guo ZV, Inagaki HK, Daie K, Druckmann S, Gerfen CR, Svoboda K (2017) Maintenance of persistent activity in a frontal thalamocortical loop. Nature. 545(7653):181–186

73. Guo ZV, Li N, Huber D, Ophir E, Gutnisky D, Ting JT, Feng G, Svoboda K (2014) Flow of cortical activity underlying a tactile decision in mice. Neuron. 81(1):179–94.

74. Gao Z, Davis C, Thomas AM, Economo MN, Abrego AM, Svoboda K, De Zeeuw CI, Li N (2018) A cortico-cerebellar loop for motor planning. Nature. 563(7729):113–116.

75. Nitz DA (2012) Spaces within spaces: rat parietal cortex neurons register position across three reference frames. Nat Neurosci. 15(10):1365–7.

76. Chen LL, Lin LH, Green EJ, Barnes CA, McNaughton BL (1994) Head-direction cells in the rat posterior cortex. I. Anatomical distribution and behavioral modulation. Exp Brain Res.101(1):8–23.

77. Jacob PY, Casali G, Spieser L, Page H, Overington D, Jeffery K (2017) An independent, landmark-dominated head-direction signal in dysgranular retrosplenial cortex. Nat Neurosci. 20(2):173–175.

78. Harvey CD, Coen P, Tank DW (2012) Choice-specific sequences in parietal cortex during a virtual-navigation decision task. Nature. 484(7392):62–8.

79. Li N, Chen TW, Guo ZV, Gerfen CR, Svoboda K (2015) A motor cortex circuit for motor planning and movement. Nature. 519(7541):51–6.

80. Inagaki HK, Inagaki M, Romani S, Svoboda K (2018) Low-Dimensional and Monotonic Preparatory Activity in Mouse Anterior Lateral Motor Cortex. J Neurosci. 38(17):4163–4185.

81. Kraus, BJ (2013) Nanconv.m.

